# Optimising tropical forest bird surveys using passive acoustic monitoring and repeated short-duration point counts

**DOI:** 10.1101/2020.08.24.263301

**Authors:** Oliver C. Metcalf, Jos Barlow, Stuart Marsden, Nárgila Gomes de Moura, Erika Berenguer, Joice Ferreira, Alexander C. Lees

## Abstract

Estimation of avian biodiversity is a cornerstone measure of ecosystem condition, with turnover in avian community composition underpinning many studies of land-use change in tropical forests. Surveys conducted using autonomous recorders have been frequently found to be more efficient than traditional point-count surveys. However, there has been limited research into optimal survey duration, despite autonomous recordings allowing for many more repeats of short-duration surveys with relative ease in comparison to traditional survey methods.

We use an acoustic dataset collected from a region of very high avian biodiversity - the eastern Brazilian Amazon - to test the effect of using short-duration surveys to increase temporal coverage without increasing total survey duration. We use this dataset to assess whether a survey protocol consisting of 240 15-second surveys at 29 locations, ‘short-duration surveys’, has an influence on resulting alpha and gamma diversity, and detection frequency, than ‘standard-duration surveys’ of four 15-minute surveys per location.

We find that repeated short-duration surveys outperform longer duration surveys in every metric considered herein, with short-duration surveys predicted to detect approximately 50% higher alpha diversity, and 10% higher gamma diversity. Short-duration surveys also detect species more often, at more survey locations. Conversely, standard-duration surveys are almost four times more likely to produce false negatives (i.e. to fail to detect species presence). Whilst there is no difference between the proportion of uncommon species detected by the two methods, when considering species detected multiple times at multiple locations, short-duration surveys detected three times more uncommon species than standard-duration surveys.

We conclude that short-duration recorded surveys should be considered the primary method for sampling the species richness of bird communities in tropical forests and is likely to be preferable to longer duration or traditional surveys in most environments.

## Introduction

Estimation of avian biodiversity is a cornerstone measure of ecosystem condition used in a wide range of ecological and conservation applications. Understanding turnover in avian community underpins many studies of land-use change in high biodiversity environments like tropical forests. However reliable detection, identification and counting of birds can be challenging in such environments (Robinson et al., 2018) where avian species richness reaches its global asymptote (Jenkins et al., 2013). It is well documented that tropical birds can be difficult to detect and count accurately, and as a consequence of low abundance, accumulating sufficient inventory completeness can be challenging (Robinson et al., 2000; Terborgh et al., 1990).

Point counts are established as a standard survey technique for obtaining measures of bird species richness, abundance and population density, particularly in forest habitats (Bibby et al., 2000). Now that affordable and reliable passive-acoustic monitoring equipment has become readily available, autonomously recorded surveys, in which automated recording units are left to record the soundscape over extended periods, are emerging as a supplement or alternative to field-conducted point counts (Shonfield and Bayne, 2017). A recent review found that recorder-based surveys detect an average of 11% more species than traditional point counts, hereafter ‘traditional surveys’, albeit often with different species composition (Darras et al., 2019). This is alongside other benefits including reduced costs, avoidance of the effects of observer presence (Hutto and Mosconi, 1984), increased standardization (Campbell and Francis, 2011) and the capacity to record for an extended duration. Whilst automated detection and classification methods are not yet widely available to analyse all of the additional data collected (Priyadarshani et al., 2018), using autonomous recorders to collect data, whilst subsequently manually detecting and identifying the species (hereafter ‘semi-automated surveys’) can still obtain significant improvements over traditional methods.

An accompanying benefit of the ability to cost-effectively record large amounts of acoustic data is the ability to vary the duration of surveys which often isn’t logistically possible if observers need to be present in the field. A larger number of short-duration traditional surveys, conducted over the same total duration as longer surveys, have been shown to give better temporal coverage and can lead to both detection of higher species richness (Fuller and Langslow, 1984; Siegel et al., 2001) and better quality data for further modelling (Dettmers et al., 1999; Lee and Marsden, 2008; Smith et al., 1998). In tropical forests, where hyper-diverse avian communities comprise a small number of abundant species and a long tail of uncommon species (Robinson et al., 2000; Terborgh et al., 1990), traditional point counts often fail to accrue enough independent repeat detections to allow modelling of rarer species, leading to knowledge deficits for species most likely to be sensitive or vulnerable to disturbance and consequently underestimating impacts of land-cover change (Robinson et al., 2018). Semi-automated surveys allow survey protocols to focus on a high number of short-duration surveys without the costs associated with multiple repeated field visits. Despite this potential benefit of semi-automated surveys, most comparisons with traditional surveys have been conducted either simultaneously or using identical sampling methods. As with traditional surveys, several recent studies comparing semi-automated surveys indicate that using a large number of short-duration surveys to increase temporal coverage allows detection of a higher number of species than standard-duration surveys over an equal total duration (Klingbeil and Willig 2015, Bayne et al., 2017, Smith et al., 2020). However, these studies were conducted in temperate forest or arid systems in regions of relatively low species richness (n=40, n=96 and n=79 species respectively), and none used a minimum survey duration of less than one minute. Additionally, Cook and Hartley (2018) found that using 10 s duration surveys increased estimation of species prevalence compared to longer duration surveys through increased independent detections. Increased detection frequency can reduce the number of false absences, something that can have a significant negative impact on the accuracy of, in particular, species distribution modelling (Gu and Swihart, 2004; Lobo et al., 2010; Phillips et al., 2009).

The detection of species richness is influenced by two factors: availability and detectability (Kéry and Schmidt, 2008). The number of species available for detection over time (e.g. the number of species close enough to the recorder to be heard), varies as species move – for instance the number of available species would be much greater if a large mixed-species flock was inside the area of detection. The detectability of each species (e.g. whether an individual of the species makes an identifiable sound during the survey) is the probability of recording the species when it is within the recording area. This is impacted by species abundance – the more individuals available for detection increases the probability of one of them vocalising, and the frequency of vocalisation, for instance screaming pihas *Lipaugus vociferans* may vocalise for 77% of the time between 06:45-17:15 (Snow, 1961), whilst variegated antpittas *Grallaria varia* have been shown to only sing only twice in 50 days (Jirinec et al., 2018). The fluctuation in the proportion of the total species pool that is both available and detectable over time means that having a higher number of surveys makes it more likely for a survey to coincide with a period in which a high proportion of the total species pool is detectable (Figure 1). In some cases, it may be possible to predict which time periods have the highest probability of detecting a higher proportion of the species pool, suggesting survey efforts should be targeted to those periods, for instance traditional point counts are often conducted in the two hours following sunrise. However, some species only vocalize within strict temporal niches, and are only detectable at certain periods - forest falcons *Micrastur spp*., for example, only reliably vocalise before and around dawn (del Hoyo et al., 2020). Other species may have habitual movements that make them only available for detection during narrow windows. Additionally, vocalizations of rare species are likely to be largely stochastic. This means that in addition to having a higher number of repetitions, survey designs with higher temporal spread should detect a higher proportion of the total species pool. This effect should be particularly pronounced in the tropics, where higher proportion of species occur naturally at low abundance, and spatial and vocalisation niches are more tightly packed (Robinson et al., 2018; Terborgh et al., 1990), leading to higher variability in the proportion of species available per survey period, and higher turnover of species between surveys. In the tropics, the regions with the highest global diversity, the benefits of increased temporal coverage should be greatest.

**Figure 1.**
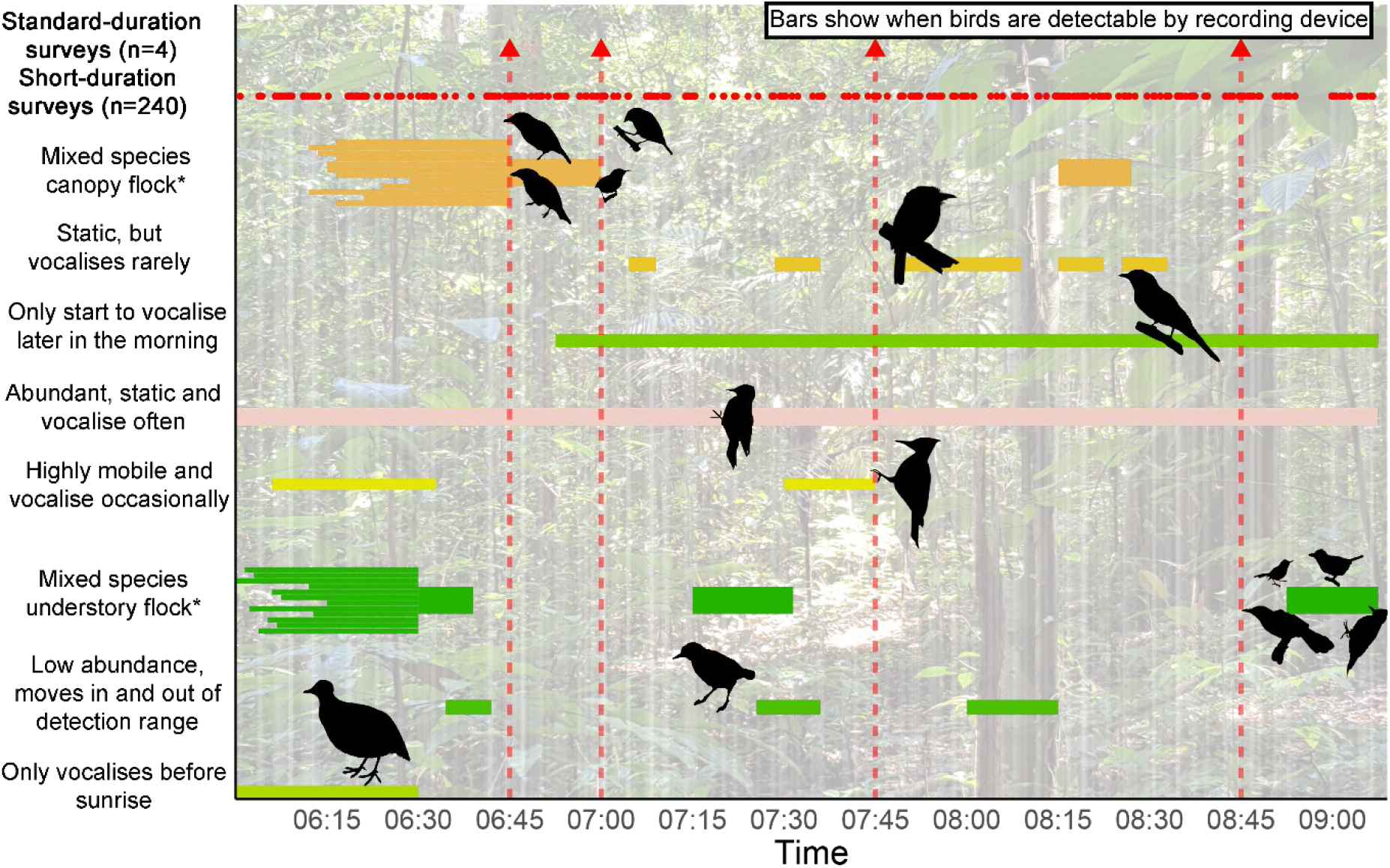
Theoretical model of long and short duration sampling regimes over one morning in the tropics. Red arrows represent four 1-minute surveys, red dots represent 1 second instantaneous surveys. The y axis shows a non-exhaustive selection of behaviours that impact detection probability. *Mixed flocks shown both prior to and after formation. Bird behaviour affecting detectability is hypothetical and not based on data.

We used an acoustic dataset collected between June and August 2018 in eastern Amazonia. We installed automated recorders across 29 transects to test the effect of using repeated short-duration surveys to increase temporal coverage, and therefore survey effectiveness, without increasing total survey duration. We compared the results with those from longer duration surveys to answer the following questions: does the use of short-duration surveys result in detection of higher species richness and a faster species accumulation? Does the use of short-duration surveys detect species more frequently and decrease the number of false absences and falsely unique occurrences? Are short-duration surveys more efficient at detecting species with low abundance?

## Methods

### Data collection

We collected acoustic data from 29 of the survey transects of the Sustainable Amazon Network (Gardner et al. 2013) located in the eastern Brazilian Amazon in the municipalities of Santarém, Belterra, and Mojuí dos Campos (latitude ~ −3.046, longitude −54.947), hereafter Santarém in the Brazilian state of Pará. Survey points were located halfway along permanent 300 m transects. All transects were located in terra firme forests and distributed across a human-disturbance gradient, comprising seven forest classes. To minimize spatial correlation, survey points were separated by a minimum distance of 2 km. Recordings were made continuously over multiple discrete time periods at each survey point, with a minimum of 13 sampling days sampled at each location. Discrete recording periods ranged in duration between 3 and 20 days. Full details of recording location, forest disturbance categories, equipment settings and recording periods for each location are given in SOM Appendix 1.

The continuous acoustic recordings were randomly and independently subsampled twice. In the first subsample (hereafter ‘standard-duration surveys’), survey periods were 15 minutes in duration, and four periods were extracted per survey point, totalling one hour of data from each transect. Across all transects, we subsampled a total of 116 standard-duration surveys. We used 15 minute survey durations as it is a commonly used point-count duration in tropical forests (Robinson et al., 2018), and as previous traditional surveys from the same location have used this survey duration (Moura et al., 2013). The second subsample (hereafter ‘short-duration surveys’) again independently sampled one hour of recordings from each survey point, but this time in the form of 240 15-second periods, totalling 6,960 surveys across all transects. We used 15-second samples for short-duration surveys as bird movement in and out of the survey point is minimized, a spectrogram of the entire survey period can be displayed at a resolution where vocalisations can be visually recognized, and as this is a trade-off between the number of complete versus truncated vocalisations which can be difficult or impossible to identify without a longer recording. All samples for both survey methods were taken in a two-and- a-half-hour period starting 30 minutes before sunrise. Subsampling was not stratified within that period, but standard-duration surveys commenced on the hour, or 15, 30 or 45 minutes past the hour, to avoid overlapping samples. Audio containing heavy rainfall was removed prior to initial sampling using the hardRain package in R (Metcalf et al., 2020).

### Analysis

The audio samples were analysed manually, through visually inspecting spectrograms and listening to the recordings. All identifiable avian vocalisations were assigned to species by a highly experienced ornithologist (NGM, for survey experience in the region see Moura et al., (2013), and Moura et al., (2016)). All vocalizations that could only be determined to family level were discarded from this study. During analysis it was apparent that 343 of the 6,960 short-duration surveys fell during periods of rain intense enough to significantly inhibit bird vocalization activity and/or detection. These were removed from consideration but not replaced, leading to an uneven sample size (see SOM Appendix 1). Consequently, for each survey point, we calculated both observed species richness and rarefied species richness for 45 minutes of sample effort to account for the uneven sampling effort across methods, using the iNext package in R, but patterns and results were similar to observed species richness, so only observed species richness is considered hereafter.

### Species Richness

We compared alpha and gamma diversity metrics between the two survey methodologies. First, we modelled species richness at each survey point using a linear mixed effect models in the lme4 package, using survey method as a fixed effect, survey point nested within forest disturbance class as a random effect, and a Gaussian error structure. We also calculated total species richness across all survey points (gamma diversity). For a repeat of this analysis including rarefied species richness, and data from traditional point-counts conducted in 2016, see SOM Appendix 2. To address whether the use of short surveys accrued species richness at a faster rate than standard surveys, we constructed sample-based species accumulation curves for each survey method, interpolating for 20 hours of survey effort using the iNext package.

### Detection Frequency

Next, we looked at whether short or standard surveys detected species more frequently. A detection is counted as a species being identified as present in a survey, (e.g. presence), not the number of times it is detected within a survey. We summed the total number of detections of each species by survey method, and compared the total number of detections for the species detected in both survey methods using a paired Mann-Whitney U test. As total detections are highly dependent on the total number of surveys and not necessarily reflective of improvements caused by greater temporal coverage, we also summed the number of survey points at which each species was detected. To assess the possible implications of higher detection rates on the data, we also calculated the number of species falsely found to be absent per survey point. A species was determined to be falsely absent if it was undetected at a location by one survey method, but detected at the same location by the converse method.

In addition, we looked at extreme cases of false absences, in which species were detected at only a single location by a survey method, but were actually present at other locations (hereafter ‘false uniqueness’), something that is likely to be highly detrimental to the accuracy of habitat modelling in particular. As most analysis of this type are directed at the habitat level we analysed this at the scale of forest class, and calculate the proportion of the total species richness of each forest class that was determined to be falsely unique species. The seven forest classes are: undisturbed forest (five survey points), selectively-logged forest (four survey points), secondary forest - forest recovering from complete historical clearance *sensu* Putz and Redford, (2010) (three points), and four categories of burnt forest. The four burnt categories were categorised dependent on whether they burnt during the extensive El Niño-induced fires in 2015 and whether they have been selectively logged, with all logging occurring prior to 2015. The categories are; burned in 2015 but never logged (five points), logged and burned prior to 2015 (four points), logged and burned in 2015 (five survey points) and logged and burned both before 2015 and in 2015 (three survey points).

### Sensitivity to abundance

To test if short surveys detected more rare species, we compared the relative abundance of species detected by both methods using chi-squared tests. We designated each species as common, fairly common, or uncommon, using the Stotz et al., (1996) species trait database. Species marked as intermediate between two abundance classes in Stotz were assumed to belong to the rarer class, categories marked as uncertain were assumed to be correct, and we combined the categories of uncommon, patchily distributed and rare. Species nomenclature was updated in line with the taxonomy of de Piacentini et al., (2015), and relative abundance data for species absent from Stotz were taken from the literature. We also tested whether short-duration surveys detected each rare species more often. To ensure that any increase in detection of rare species was not caused by repeatedly detecting a single individual more often with short-duration surveys, we also compared the number and proportion of species that were detected from a minimum of two transects and with >10 total detections (hereafter ‘multiple detections’).

## Results

### Species Richness

We detected higher alpha and gamma diversity (Figure 2A) using short surveys. In total, we detected 245 species; 224 species using short surveys with a median of 4.0±0.02 (SE) species and 204 species using standard surveys with a median of 19.5±0.68 species per survey. The linear mixed effects model predicted that short surveys detect 22.9 species more per survey point than standard surveys, with short surveys detecting 66.27±3.77 (SE) per point and standard surveys detecting 43.37±3.77 species per point. Short surveys detected 41 species not detected by standard surveys across the landscape, whilst standard surveys detected 21 species not detected by short surveys.

**Figure 2:**
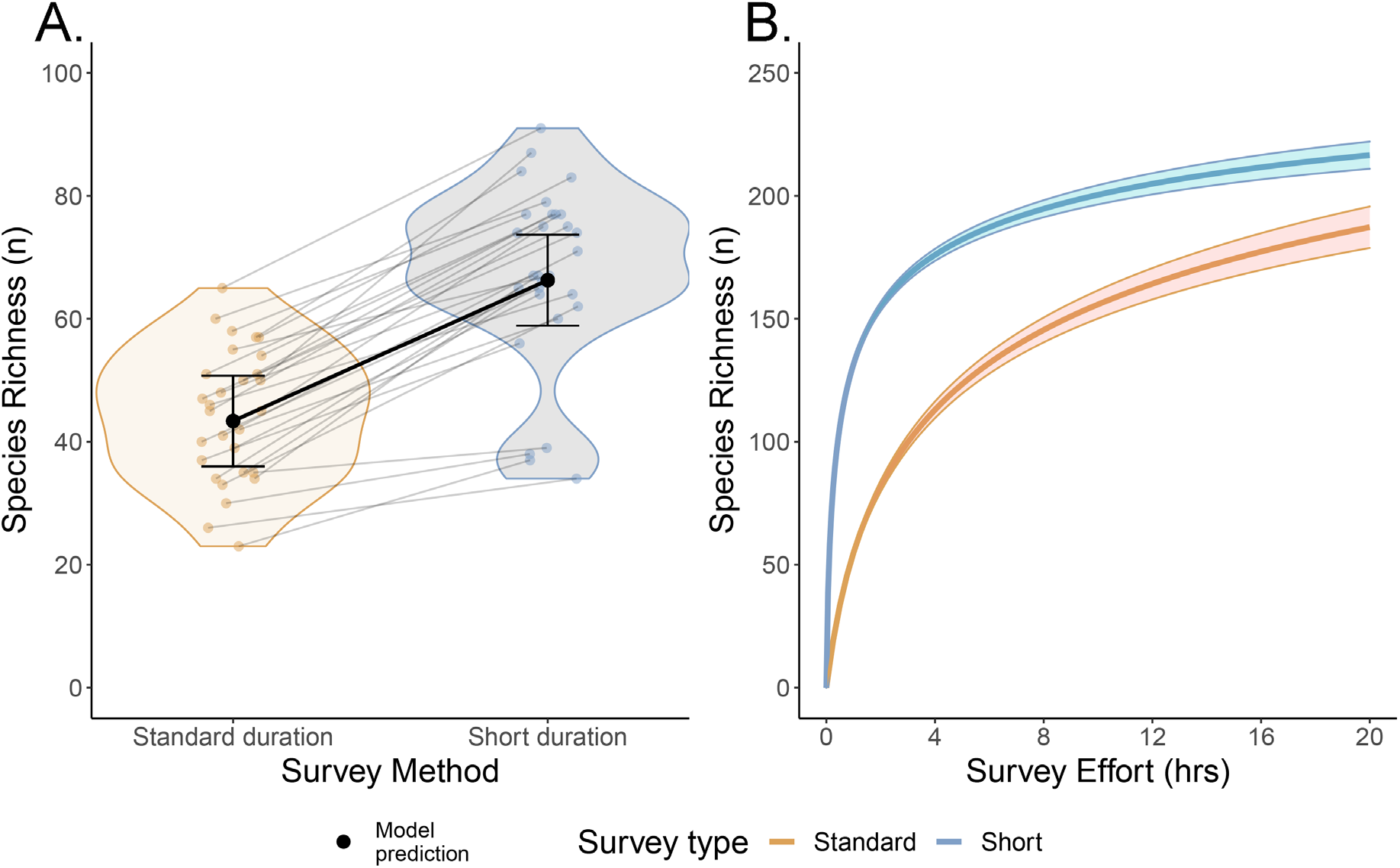
A; Comparison of the species richness detected at each of 29 survey points when using standard surveys, comprised of four 15-min periods, or short-duration surveys of 240 15-s periods. 1B; Sample-based species accumulation curves for the two survey methods, showing interpolated predictions up to 20 hours survey effort.

We found that for sample-based rarefaction/extrapolation by survey method (Fig. 2B), short-duration surveys led to steep increases in species accumulation up to around four hours of survey effort, with 176 ± 2 (SE) species detected, and then stabilized with slower species accumulation continuing up to 20 hours. In contrast, standard surveys showed a shallower curve, in which the accumulation did not seem to stabilize. Standard surveys detect lower species richness at all quantities of survey effort and were predicted to detect 187 ± 8 (SE) species after 20 hours of survey effort, compared to 217 ± 5 species by short surveys. Short surveys were predicted to take just 11 hrs 23 mins to achieve the same species total as standard surveys did in all surveys (204 species, 29 hrs).

### Detection Frequency

Detection frequency also significantly increased with short-duration surveys. Species were detected more often, a median of 47 ± 18.9 (SE) times compared to just 7 ± 1.0 for standard surveys (V=15865, p<0.001), and at more transects, 8 ± 0.57 to 4 ± 0.47 (V=976, p<0.001) (Fig 3A). Additionally, standard surveys detected 65% of all species fewer than ten times, and only six species were detected more than 50 times, with a maximum of 64 detections for grey antbird *Cercomacra cinarescens*. Short-duration surveys detected only 33% of all species fewer than ten times, recorded 40% of all species more than 50 times, recorded three species more than 1,000 times and had a maximum of 1,821 detections for bright-rumped attila *Attila spadiceus*.

**Figure 3.**
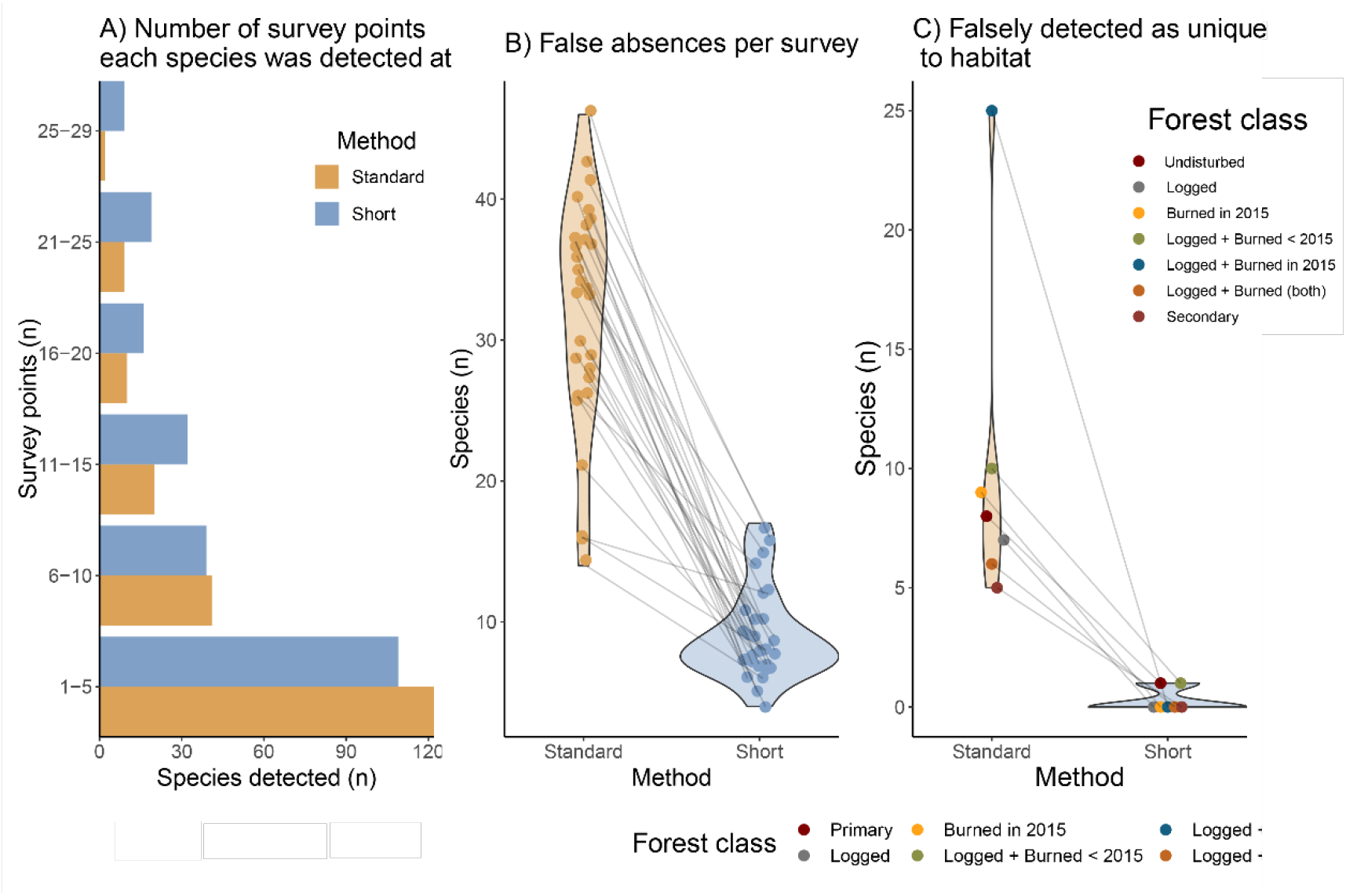
Frequency of detection. A; The number of survey points each species was detected according to survey method; short-duration surveys (240 x 15 s per survey point) and standard-duration surveys (116 x 15 min). B; The number of species falsely identified as absent per survey point. C; the number of species wrongly identified as unique to each forest class.

We found that the higher detection frequency had a striking effect on the accuracy of species absences with standard-duration surveys producing 927 false absences compared to just 263 for short-duration surveys. Every survey point had fewer false absences with short-duration surveys than standard-duration (Fig. 3B), and at one location, 50 species were detected with standard duration surveys, but a further 46 were missed – whilst only nine were missed with short-duration surveys.

This pattern was also apparent when looking at species that were only detected in a single forest class, but were actually present in others as well. There were only two species which short surveys wrongly identified as unique to a forest class, compared to 70 by standard-duration surveys. One forest class, logged and burned in 2015, had an exceptionally high error rate using standard-duration surveys, with 25 species or 21% of the total detected species at that class being wrongly detected as unique - something that could be highly misleading in habitat or distribution modelling. As the other forest class that was also logged and burnt had an almost equally high error rate, it seems likely that the two classes shared a high proportion of species at low abundance that were not well detected by standard-duration surveys, but were by short-duration surveys.

### Sensitivity to abundance

Short-duration surveys detected a mean 10%±0.7(SD) more species for common, fairly common and uncommon birds. However, both survey methods detected a remarkably similar proportion of each category of relative abundance (Fig. 4). When only considering multiple detections of species (10+ total detections and detected at two or more locations), short-duration surveys detected substantially more species than standard-duration surveys, with the largest difference being for uncommon species for which short-duration surveys detected nearly three times as many species (n = 13 and 38, respectively). Furthermore, the number of uncommon species detected as a proportion of all species detected multiple times declined for standard surveys (28% to 18%) but stayed relatively stable for short surveys (29% to 25%). When analysing only standard surveys, the proportion of uncommon species in the total species pool declined from 28% for all species detected, to 18% when considering only multiple detections. For short surveys, the detection of uncommon species remained similar, regardless of the abundance metric used (29% to 25%).

**Figure 4.**
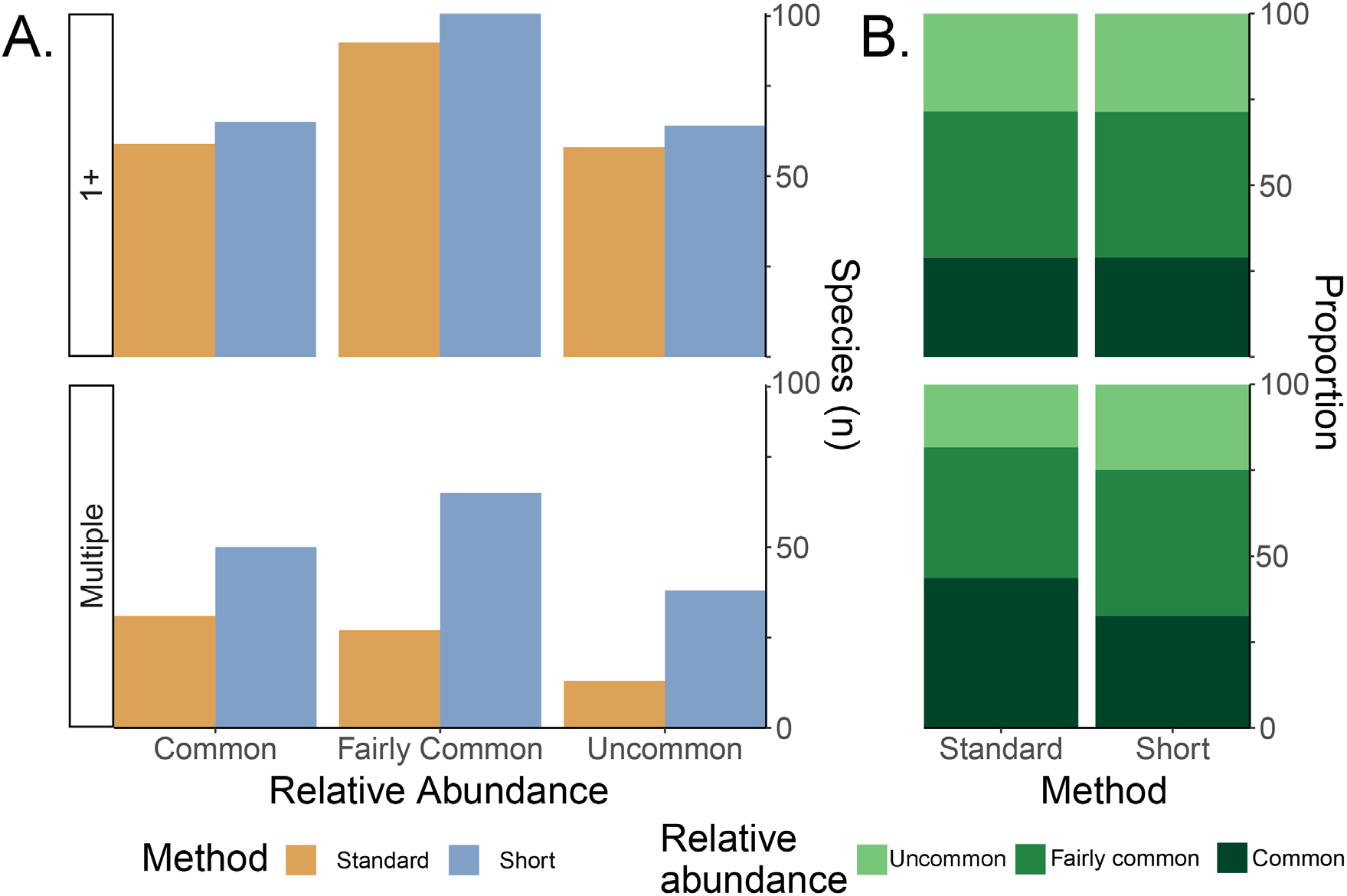
The proportion of common, fairly common and uncommon species detected using both the standard and the short survey methods.

## Discussion

We found that short-duration avian surveys using passive acoustic monitoring outperformed longer duration surveys in every metric considered, often by a substantial margin. This is particularly true for species richness, where we predict short surveys to record just over a third more species at each location, as well as finding substantially higher gamma diversity. Looking beyond species richness, short-duration surveys are also a more reliable method to obtain data for distribution and occupancy modelling. Short surveys produce far fewer false negatives for the presence of species, and identify far fewer species as unique to forest class, both of which can be significant hindrances in habitat and distribution modelling (Gu and Swihart, 2004; Kramer-Schadt et al., 2013). The ability to reliably repeatedly detect rarer species also means that short-duration surveys are more robust to low relative abundance, which can be advantageous in surveying bird communities, particularly in the tropics (Robinson and Curtis, 2020).

We have not conducted sensitivity analysis to optimise the duration of surveys. However, one previous study compared survey durations of ten, five, three, two and one minute across equivalent cumulative periods and found species detection rates increased as survey durations decreased (Bayne et al., 2017). This, alongside our own results suggest that by shortening survey duration and increasing the temporal spread of surveys, species accumulation will continue to increase. In fact, whilst estimates of abundance from acoustic surveys remain in their infancy due to difficulty with estimating distance from audio data (Darras et al., 2016; Yip et al., 2017), using near instantaneous survey durations could resolve the issue of movement in and out of detection range during the survey period. However, there are some inhibiting factors to suggest that extremely small durations (<10 s) may not be beneficial overall. Firstly, NGM reported issues with identification of calls with the shorter duration surveys due to vocalisations being truncated at the start and end of the recordings, or absence of patterns in vocalisations that can be important cues in longer recordings. Secondly, and potentially more significantly, there are substantially higher analysis costs associated with short-duration surveys due to the need to record results repeatedly, in this case all of the meta-data and species presence data associated with a survey would need to be recorded 60 times for 15 second recordings, compared to just once for 15 minute recordings. Whilst the extra time required in analysis is undoubtedly substantial, it could be offset by the use of specialist software(e.g. BORIS (Friard and Gamba, 2016)), and by lower total survey duration required due to the increased species accumulation rate.

## Conclusion

We believe that short-duration recorded surveys should be considered the standard and primary method for sampling bird communities in tropical forests. There is strong evidence that surveys conducted on longer autonomous sound recordings outperform human observations when surveying birds (Darras et al., 2019), suggesting that autonomous surveys should be used preferentially or in combination with traditional point-count surveys. Given the additional benefits of short-duration surveys, we believe that within tropical forest environments manually conducted point counts should mainly be employed as a supplement to short-duration surveys. Exceptions include when autonomous recordings are not possible, for example if equipment cost is too high, when estimates of abundance are of higher priority than estimates of species richness, and when a high proportion of non-vocalising species are expected. Longer duration autonomous surveys offer little benefit over either human observations or short-duration surveys, except when analysis time is of high priority. Whilst a combination of traditional and autonomous survey techniques should still be considered the gold standard for conducting bird species inventories (Robinson and Curtis, 2020), if only a single survey method is to be used, repeated short-duration surveys are likely to be the most effective.

## Acknowledgements

We thank the Large Scale Biosphere-Atmosphere Program (LBA) for logistical and infrastructure support during field measurements. We are very grateful for the hard work of our field and laboratory assistants: Marcos Oliveira, Gilson Oliveira, Renílson Freitas, Josué Jesus de Oliveira, and Amanda Cardoso. We are also grateful to Liana Rossi and Filipe França for logistical field support in Brazil, and to Jack Shutt for advice on statistical analysis. Fieldwork in Brazil was supported by research grants ECOFOR (NE/K016431/1), and AFIRE (NE/P004512/1), PELD-RAS (CNPq/CAPES/PELD 441659/2016-0). The authors have no conflicts of interest to declare.

## Author’s contributions

xxx

## Data Availability

We will make the species presence data (per survey and survey method) available in the Dryad data repository on acceptance of this article.

## Notes

### Competing Interest Statement

The authors have declared no competing interest.

