## Supplementary Online Materials for "Optimising tropical forest bird surveys using passive acoustic monitoring and repeated short-duration point counts"

Appendix 1: Survey locations, total effort and number of samples

| Point | Transect name | Latitude | Longitude | First recording date | Last recording date | Duration (mins) | Habitat type (primary unless stated) | Short-duration surveys (n) | Standard-duration surveys (n) |
|---|---|---|---|---|---|---|---|---|---|
| 1 | B112 T12 | -2.693 | -54.452 | 06/07/2018 | 30/07/2018 | 22992 | Logged and burned in 2015 | 231 | 4 |
| 2 | B112 T8 | -2.693 | -54.495 | 06/07/2018 | 31/07/2018 | 26397 | Logged | 231 | 4 |
| 3 | B125 T9 | -2.804 | -54.568 | 02/08/2018 | 16/08/2018 | 19716 | Logged and burned in 2015 | 239 | 4 |
| 4 | B129 T10 | -2.726 | -54.777 | 30/06/2018 | 16/08/2018 | 17862 | Logged and burned < 2015 | 232 | 4 |
| 5 | B129 T11 | -2.706 | -54.786 | 30/06/2018 | 13/08/2018 | 21970 | Secondary | 214 | 4 |
| 6 | B129 T5 | -2.714 | -54.749 | 30/06/2018 | 13/08/2018 | 21700 | Logged and burned < 2015 | 238 | 4 |
| 7 | B160 T10 | -2.819 | -54.882 | 14/06/2010 | 28/06/2018 | 19333 | Logged | 192 | 4 |
| 8 | B260 T1 | -3.002 | -54.895 | 14/06/2018 | 28/06/2018 | 19145 | Logged and burned in 2015 | 231 | 4 |
| 9 | B260 T4 | -3.02 | -54.857 | 14/06/2018 | 12/07/2018 | 25897 | Logged and burned in 2015 | 224 | 4 |
| 10 | B260 T5 | -2.984 | -54.878 | 14/06/2018 | 28/06/2018 | 19081 | Logged and burned – both | 230 | 4 |

| | | | | | | | | | |
|---|---|---|---|---|---|---|---|---|---|
| 11 | B260 T6 | -3.00 | -54.877 | 14/06/2018 | 28/06/2018 | 19113 | Logged | 219 | 4 |
| 12 | B261 T10 | -3.018 | -55.005 | 12/06/2018 | 05/07/2018 | 30769 | Burned in 2015 | 222 | 4 |
| 13 | B261 T8 | -3.029 | -55.011 | 12/06/2018 | 05/07/2018 | 31468 | Burned in 2015 | 227 | 4 |
| 14 | B261 T9 | -3.04 | -55.015 | 12/06/2018 | 13/08/2018 | 28212 | Burned in 2015 | 218 | 4 |
| 15 | B307 T3 | -3.13 | -54.857 | 20/07/2018 | 03/08/2018 | 27146 | Logged and burned – both | 239 | 4 |
| 16 | B307 T7 | -3.147 | -54.838 | 20/07/2018 | 03/08/2018 | 18932 | Logged and burned – both | 240 | 4 |
| 17 | B357 T4 | -3.283 | -54.854 | 18/06/2018 | 02/07/2018 | 19371 | Secondary | 210 | 4 |
| 18 | B363 T3 | -3.296 | -54.963 | 02/07/2018 | 23/07/2018 | 18958 | Undisturbed | 239 | 4 |
| 19 | B363 T5 | -3.336 | -54.984 | 19/07/2018 | 10/08/2018 | 17827 | Undisturbed | 230 | 4 |
| 19 | B363 T6 | -3.336 | -54.956 | 03/07/2018 | 15/08/2018 | 18958 | Undisturbed | 240 | 4 |
| 20 | B363 T7 | -3.32 | -54.96 | 21/06/2018 | 15/08/2018 | 26246 | Undisturbed | 239 | 4 |
| 21 | B363 T8 | -3.329 | -54.972 | 21/06/2018 | 15/08/2018 | 26119 | Undisturbed | 239 | 4 |
| 22 | B399 T7 | -3.429 | -54.843 | 14/07/2018 | 31/07/2018 | 22085 | Logged | 240 | 4 |
| 23 | B399 T8 | -3.482 | -54.862 | 14/07/2018 | 10/08/2018 | 25334 | Secondary | 240 | 4 |
| 24 | B399 T10 | -3.464 | -54.907 | 14/07/2018 | 31/07/2018 | 21897 | Logged and burned < 2015 | 237 | 4 |

| | | | | | | | | | |
|---|---|---|---|---|---|---|---|---|---|
| 25 | B69 T11 | -2.574 | -54.671 | 13/06/2018 | 29/06/2018 | 25989 | Logged and burned < 2015 | 199 | 4 |
| 26 | B69 T8 | -2.515 | -54.675 | 13/06/2018 | 29/06/2018 | 26103 | Logged and burned in 2015 | 225 | 4 |
| 27 | BExtra T2 | -2.938 | -54.988 | 13/06/2018 | 29/06/2018 | 25687 | Burned in 2015 | 219 | 4 |
| 28 | BExtra T3 | -2.928 | -55.002 | 19/06/2018 | 09/07/2018 | 32277 | Burned in 2015 | 233 | 4 |

Transect name: B=catchment number, T= transect number as detailed in Gardner et al.,(2013)

Appendix 2: Manual surveys

We compared this acoustic data with traditional point count surveys conducted at the same sites by three experienced observers (ACL, NGM and Sidnei Dantas, see Moura et al., (2013), Moura et al., (2016), Henriques et al., (2003)) between 15-26$^{th}$ November 2016. Surveys lasted 15 minutes each, and were conducted between 05:45 and 09:45, and two surveys were conducted from each of 0 m, 150 m and 300 m along the transect, so that each transect was surveyed six times in total. The detection method was noted for each species per survey as either visual or auditory. In all other respects, the surveys followed the protocols set out in previous published surveys at the site (Lees et al., 2013). There are distinct differences between the traditional surveys and the recorded surveys; being conducted at the start of the rainy season when birds are expected to be most vocally active and hence easily detected, across a greater spatial scale (three points, at 0m, 150 m and 300m along each transect), for 90 minutes rather than 60 at each transect, and over a slightly longer survey window each morning. Furthermore, ten of the transects were fire-damaged during El Nino events in 2015, and therefore have significantly different vegetation structure between 2016 and now owing to two years worth of post-fire regeneration (Berenguer et al., 2018), whilst the structure of the other transects remains similar.

We modelled the resulting species richness scores using linear mixed effects models in the lme4 package, using survey method and presence of fire in 2015 as fixed effects and transect nested within disturbance class as a random effect.

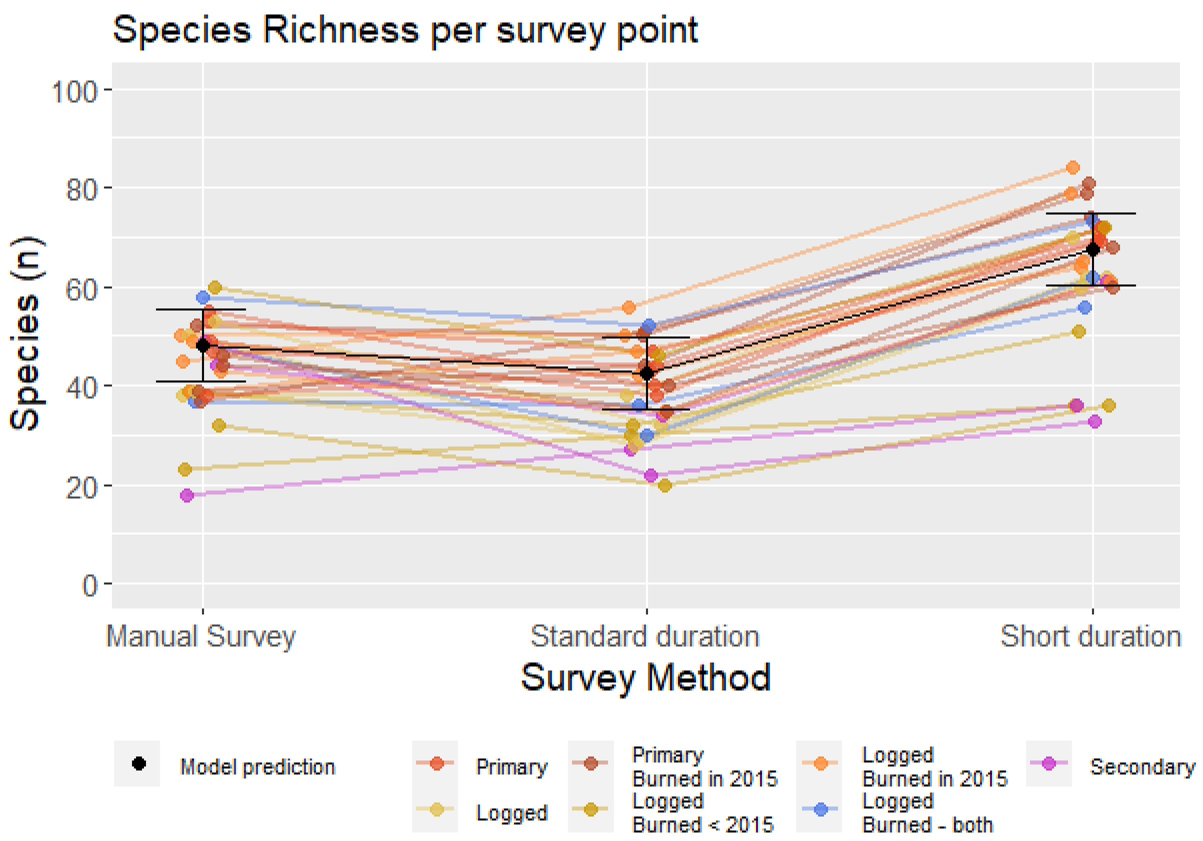


Traditional surveys detected a total of 255 species, higher than either of the other methods. However, traditional surveys detected 48±3.7 (SE) species per transect, which is 5.4±1.85 more than standard-duration surveys, but 19.4±1.85 species less than short-duration surveys.

Whilst we statistically account for the differences in survey method where possible using rarefied species richness data (Oksanen et al., 2019), we believe that the traditional surveys are detecting higher species richness than had they been conducted in the same season of 2018 using identical protocols as the autonomously recorded surveys – particularly given the results of Darras et al., (2019). However, we have included the comparison here as we believe it is useful to readers to place the results of the comparison between short and long duration surveys with autonomous audio recordings in the context of the regional species pool likely to be detected by traditional point counts. It further emphasises the efficacy of short-duration surveys that despite the considerable biases in favour of the traditional surveys, short-duration recorded surveys still recorded substantially higher species richness.

Appendix 3: Model Specifications and Diagnostic Plots

**Model 1**

Linear mixed model fit by REML ['lmerMod']

Formula: Species.Richness ~ method + (1 | ForestClass) + (1 | SurveyPoint)

Scaled residuals:

| Min | 1Q | Median | 3Q | Max |
|---|---|---|---|---|
| -1.9599 | -0.5124 | 0.1788 | 0.5314 | 1.4423 |

Random effects:

| Groups Name | Variance | Standard Deviation |
|---|---|---|
| SurveyPoint (Intercept) | 71.54 | 8.458 |
| ForestClass (Intercept) | 73.46 | 8.571 |
| Residual | 33.19 | 5.761 |

Number of obs: 58, groups: SurveyPoint, 29; ForestClass, 7

Fixed effects:

| Groups Name | Estimate | Std. Error | t value |
|---|---|---|---|
| (Intercept) | 43.378 | 3.770 | 11.51 |
| methodShortDuration | 22.897 | 1.513 | 15.13 |

Correlation of Fixed Effects: methodShortDuration -0.201

Shapiro-Wilk normality test: W = 0.96942, p-value = 0.1503

Levene's Test for Homogeneity of Variance (center = median)

Df F value Pr(>F)

group 1 1.1896 0.2801

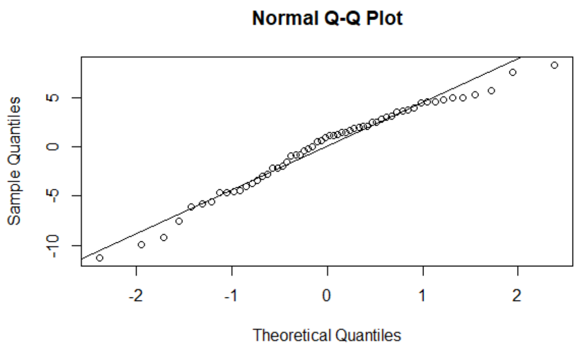


*Figure 1: Model 1 Q-QPlot*

| **Model 2** | | | | |
|---|---|---|---|---|
| Generalized linear mixed model fit by maximum likelihood (Laplace Approximation) [glmerMod]<br><br>Family: poisson ( log ) | | | | |
| Formula: Species ~ Method + (1 \| ForestClass) + (1 \| SurveyPoint) | | | | |
| Scaled residuals: | | | | |
| AIC<br><br>357.8018 | BIC<br><br>366.0436 | logLik<br><br>-174.9009 | deviance<br><br>349.8018 | df.resid<br><br>54 |
| Random effects: | | | | |
| Groups Name | Standard Deviation | | | |
| SurveyPoint (Intercept) | 0.1114 | | | |
| ForestClass (Intercept) | 0.1303 | | | |

| | |
|---|---|
| Residual | 33.19 |
| Number of obs: 58, groups: SurveyPoint, 29; ForestClass, 7 | |
| Fixed effects: | |
| (Intercept) 3.44 | MethodStandardDuration -1.26 |
| Shapiro-Wilk normality test: W = 0.98295, p-value = 0.588 | |
| The square root of the scale parameter: 1.029843 | |
| Levene's Test for Homogeneity of Variance (center = median)<br>Df F value Pr(>F)<br>group 1 0.0129 0.9101 | |

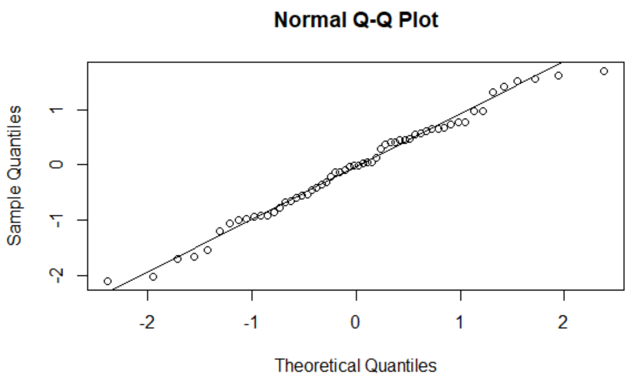


*Figure 2: Model 2 Q-Q Plot*
